# Praja1 ubiquitin ligase facilitates degradation of polyglutamine proteins and suppresses polyglutamine-mediated toxicity

**DOI:** 10.1101/2020.12.13.417964

**Authors:** Baijayanti Ghosh, Susnata Karmakar, Mohit Prasad, Atin K. Mandal

**Author notes:** Address correspondence to Atin K. Mandal, Atin K. Mandal, Division of Molecular Medicine Bose Institute, P-1/12 C.I.T. Scheme VIIM, Kolkata, India, 700054.

## Abstract

A network of chaperones and ubiquitin ligases sustain intracellular proteostasis, and is integral in preventing aggregation of misfolded proteins associated with various neurodegenerative diseases. Using cell-based studies of polyglutamine (polyQ) diseases: Spinocerebellar ataxia Type 3 (SCA3) and Huntington’s disease (HD), we aimed to identify crucial ubiquitin ligases that protect against polyQ aggregation. We report here that Praja1 (PJA1), a Ring-H2 ubiquitin ligase abundantly expressed in the brain is diminished when polyQ repeat proteins (Ataxin-3/Huntingtin) are expressed in cells. PJA1 interacts with polyQ proteins and enhances their degradation resulting in reduced aggregate formation. Down-regulation of PJA1 in neuronal cells increases polyQ protein levels vis-a-vis their aggregates rendering the cells vulnerable to cytotoxic stress. Finally, PJA1 suppresses polyQ toxicity in yeast and rescues eye degeneration in transgenic *Drosophila* model of SCA3. Thus, our findings establish PJA1 as a robust ubiquitin ligase of polyQ proteins and induction of which might serve as an alternative therapeutic strategy in handling cytotoxic polyglutamine aggregates.

## INTRODUCTION

Polyglutamine (PolyQ) diseases are neurodegenerative disorders arising from CAG trinucleotide repeat expansion in the protein coding region of the respective disease gene. Currently, nine such polyQ disorders have been identified: Spinocerebellar Ataxia (SCA) Type 1, 2, 3, 6, 7, and 17; Huntington’s disease (HD); Dentatorubral Pallidoluysian atrophy (DRPLA); and Spinal and Bulbar Muscular Atrophy (SBMA) which are progressive and dominantly inherited (except SBMA) (Zoghbi and Orr, 2000; Stoyas and La Spada, 2018). The most common among these neurodegenerative disorders with distinct pathological and clinical attributes: Spinocerebellar Ataxia Type 3 (SCA3) and Huntington’s disease (HD) are characterised by formation of amyloid like toxic intracellular aggregates, which are hallmarks of all polyQ diseases. These insoluble intra-neuronal aggregates and/or inclusions are formed at the cerebellar Purkinje neurons, brain stem and spinocerebellar tracts, and are caused as a result of a gain-of-function effect of polyglutamine expansion of the ataxin-3/huntingtin proteins causing neuronal dysfunction and death (Zoghbi and Orr, 2000; Paulson, 2012; Paulson *et al.*, 2017). These inclusions are immunoreactive for ubiquitin and sequester components of ubiquitin proteasome system (UPS), molecular chaperones and several transcription factors (Suhr *et al.*, 2001; Waelter *et al.*, 2001). The association of polyQ aggregates with the components of the protein quality control (PQC) machinery induces cellular response to manage the accumulation of misfolded proteins either by refolding them by molecular chaperones or degrading them by the degradation machinery. Multiple lines of evidence have demonstrated the function of selective molecular chaperones, the proteasome system and autophagy in suppression of aggregate formation and modulation of polyQ pathogenesis (Chai *et al.*, 1999; Brehme *et al.*, 2014; Djajadikerta *et al.*, 2019).

Cellular proteostasis is finely orchestrated by the two arms of PQC system: molecular chaperones and degradation machinery. Apart from their role in protein folding, chaperones hand over misfolded proteins to ubiquitin ligases for tagging them with ubiquitin for their degradation via the proteasome or autophagic pathway. The presence of approximately 600 ubiquitin ligases in the mammalian system (Deshaies and Joazeiro, 2009) suggests a cohort of ubiquitin ligases labouring in unison to protect cells from accumulation of misfolded proteins and their aggregates and subsequent cytotoxic stress (Theodoraki *et al.*, 2012; Kuang *et al.*, 2013). PolyQ expanded ataxin-3 and huntingtin are similarly degraded by multiple ubiquitin ligases, such as Carboxy Terminus of Hsp70-Interacting Protein (CHIP) which promotes degradation of polyQ proteins and reduces their aggregation in cell and animal models of SCA3 and HD (Jana *et al.*, 2005; Miller *et al.*, 2005). ER-associated E3 ubiquitin ligase Autocrine Motility Factor Receptor (AMFR) or Gp78 and E6-Associated Protein (E6-AP) provide cytoprotection against mutant SOD1 and mutant ataxin-3 mediated toxicity (Ying *et al.*, 2009; Mishra *et al.*, 2013). Parkin is impaired in Parkinson’s disease, and interacts with expanded polyglutamine proteins and promotes their degradation (Tsai *et al.*, 2003). Recent studies also establish the role of Mahogunin RING finger −1 (MGRN1) and Itchy E3 ubiquitin ligase (ITCH) in promoting the degradation of expanded polyglutamine proteins and suppression of polyQ mediated toxicity (Chhangani and Mishra, 2013; Chhangani *et al.*, 2014). Since polyQ disorders are age-onset neurodegenerative disorders where the cellular machinery eventually loses the robustness and efficiency of the PQC, stimulating the components of the PQC has been commonly used as a therapeutic strategy to ameliorate the pathogenesis of these disorders. Interestingly, ataxin-3 is a deubiquitinating enzyme (DUB) that tightly regulates the activity of CHIP and Parkin, E3 partners of ataxin-3 (Durcan and Fon, 2011; Durcan *et al.*, 2011; Scaglione *et al.*, 2011; Durcan and Fon, 2013). Moreover, reduced levels of CHIP and Parkin in mouse models of SCA3 imply the involvement of other robust E3 ligases restricting disease pathogenesis (Durcan and Fon, 2011; Durcan *et al.*, 2011; Scaglione *et al.*, 2011).

Analysing the interactome of ataxin-3 protein by String 9.0 and BIOGRID databases, we came across a particular ubiquitin ligase Praja1 (PJA1), a RING-H2 ubiquitin ligase which had predictive interaction with ataxin-3, but its function was unknown. PJA1 is expressed ubiquitously in various tissues, although the highest expression is observed in the brain. Interestingly, the PJA1 gene is located in a specific region of the X-chromosome, which is associated with numerous X-linked cognitive disorders and a contiguous gene deletion results in craniofrontonasal disorder (Mishra *et al.*, 1997; Wieland *et al.*, 2007). Moreover, a loss-of-function PJA1 variant was linked to neurodevelopmental disorders associated with epilepsy or/and craniofacial abnormalities (Suzuki *et al.*, 2020). Notably, among 29 other genes (linked to neurodegenerative disorders) PJA1 gene is downregulated in PR5 mutant Tau transgenic mice (Alzheimer’s disease) amygdala (Ke *et al.*, 2012). In several other studies, PJA1 has been shown to be differentially regulated in Huntington’s disease (HD) mouse model, amyloid precursor protein (APP) treated organotypic hipppocampal slice cultures, bipolar disorder affected orbitofrontal cortex (Stein *et al.*, 2004; Cui *et al.*, 2006; Ryan *et al.*, 2006). In a recent study, PJA1 has also been shown to suppress cytoplasmic TDP-43 inclusions (Watabe *et al.*, 2020).

We report here that PJA1 acts as an ubiquitin ligase of polyQ proteins, ataxin-3 and huntingtin, and inhibits their accumulation as toxic aggregates. Thereby, PJA1 suppresses polyQ mediated toxicity in yeast and *Drosophila* SCA3 transgenic model. Additionally, PJA1 silencing in mouse neuronal cells shows marked increase in SCA3 and HD aggregates, indicating crucial role of PJA1 in disease pathogenesis when its function is compromised at old age. Thus, we demonstrate PJA1’s function as a critical ubiquitin ligase present in brain, the levels of which can be manipulated to strategise a potential therapy against polyQ disorders in general.

## RESULTS

### PJA1 is dysregulated by polyQ proteins, ataxin-3 and huntingtin

Reduced efficiency of protein quality control machinery leads to development of age onset neurodegenerative disorders which result in accumulation of terminally misfolded toxic proteins (Bence *et al.*, 2001; Bennett *et al.*, 2005; Thibaudeau *et al.*, 2018). For instance, ubiquitin ligase *viz.* MGRN1, CHIP, Parkin levels and their functions are compromised in neurodegenerative diseases including polyQ disorders such as SCA3 and HD (Durcan and Fon, 2011; Durcan *et al.*, 2011; Scaglione *et al.*, 2011; Chhangani and Mishra, 2013; Durcan and Fon, 2013). Since Praja1 ubiquitin ligase is highly enriched in brain tissue including regions of the cerebellum, cerebral cortex, medulla, occipital lobe, frontal, temporal lobe and the putamen (Yu *et al.*, 2002), we speculated that its level might be reduced in neurodegeneration and result in accumulation of misfolded polyQ proteins. To check our hypothesis we examined the level of endogenous Praja1 (PJA1) upon over-expression of polyQ expanded proteins ataxin-3 and huntingtin in mouse neuroblastoma Neuro2A (N2A) cells. Truncated 20Q and 80Q ataxin-3 in pEGFP-N1 (henceforth 20QT and 80QT) and N-terminal fragment of HTT 16Q in pEGFP-C1 and 83Q in pDsRed (16Q HTT and 83Q HTT) vector and the corresponding empty vectors were transfected in Neuro2A cells and PJA1 transcripts were analysed. Interestingly, the mRNA levels of PJA1 in both ataxin-3 (20QT and 80QT) and huntingtin (16Q and 83Q) over-expressed N2A cells were decreased by approximately 0.6 fold and 0.5 fold respectively as compared to control cells (Figure 1, A and C). However, no reduction of PJA1 transcripts was noticed in case of the empty vectors. The reduction in the mRNA transcripts was also reflected in the endogenous protein levels of PJA1 in both ataxin-3 and huntingtin over-expressed N2A cells (Figure 1, B and D). The diminution of PJA1 levels during polyglutamine disorders is in coherence with the decline in its levels in PR5 mutant Tau transgenic mice amygdala, bipolar disorder affected orbitofrontal cortex and other cognitive disorders (Stein *et al.*, 2004; Cui *et al.*, 2006; Ryan *et al.*, 2006; Wieland *et al.*, 2007).

**FIGURE 1.**
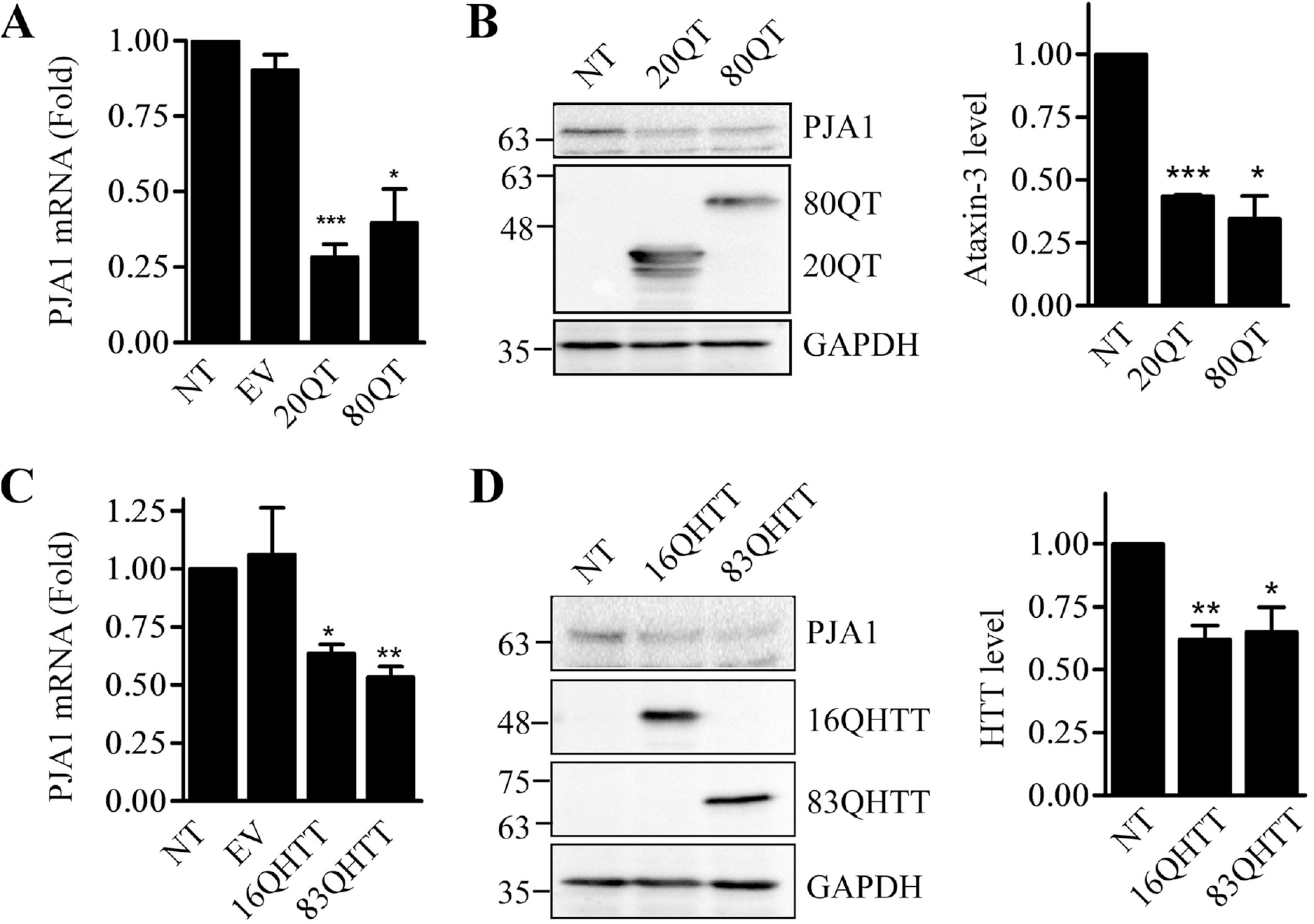
PJA1 is dysregulated by polyQ proteins. (A and C) PJA1 transcript level was decreased upon over-expression of ataxin-3 and huntingtin protein. Normal and polyglutamine (polyQ) expanded ataxin-3 (A) and huntingtin (C) plasmids and their corresponding empty vectors pEGFP-C1 and pDsRED-C1 were transfected in mouse neuroblastoma, Neuro2A cells. 72 hours after transfection, total RNA was isolated from the cells and subjected to quantitative real-time PCR. Data were collected from three separate experiments and normalised to the levels of GAPDH. Values are the mean ± S.D. Significance of the data was calculated by t-test. p <0.05 for *, p<0.01 for **, p<0.001 for *** compared to non-transfected cells. (B and D) PJA1 protein level is reduced in ataxin-3 and huntingtin transformed cells. The lysates from (A) and (C) over-expressing ataxin-3 and huntingtin respectively were immunoblotted with PJA1 and GAPDH antibody. Lysate shows the expression of ataxin-3 and huntingtin protein by GFP and DsRed antibody. The bands were quantified, normalized with GAPDH and plotted as bar diagram. Error bar represents SD of three independent experiments. Significance was calculated by t-test, *(p<0.05), **(p<0.01), *** (p<0.001). NT stands for non-transfected cells.

### PJA1 interacts with normal and polyQ expanded proteins and colocalizes with polyQ aggregates

Since overexpression of polyQ expanded proteins leads to a marked reduction in PJA1 levels we cogitated a physical interaction between PJA1 ubiquitin ligase and polyQ proteins. Intracellular aggregates tend to sequester several proteins including chaperones, ubiquitin ligases, proteasomal subunits etc (Stenoien *et al.*, 1999). Hence, we inspected whether PJA1 was recruited to the aggregates formed by the polyQ proteins. To investigate that, DsRed-PJA1 was co-transfected with GFP-tagged ataxin-3 constructs in HEK293T cells. Expression of polyQ expanded ataxin-3 (80QT and 130QF) shows large cytoplasmic aggregates and these aggregates indubitably associated with PJA1 as observed by confocal microscopy (Figure 2, A and B). However, small aggregates of 20QT ataxin-3 were observed in very few cells, which also colocalized with PJA1 (Figure 2C). Likewise, polyQ expanded huntingtin (83Q HTT) aggregates colocalized with PJA1 (Figure 2Di).

**FIGURE 2.**
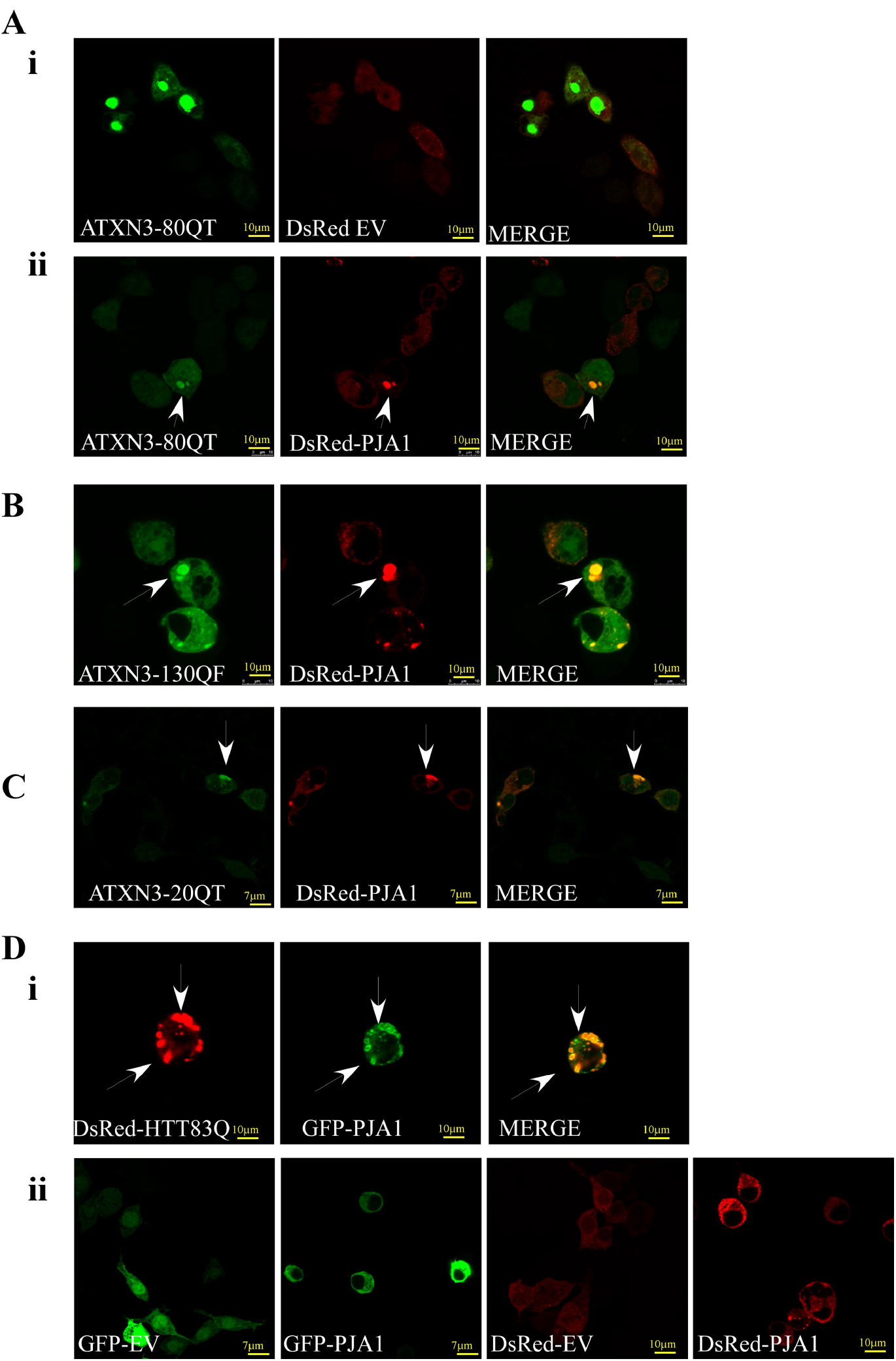
PJA1 colocalizes with polyQ aggregates. (A-C) Co-localization of PJA1 with ataxin-3. PJA1 cloned in pDsRed vector was co-transfected with 80QT (Aii), 130QF (B) and 20QT (C) ataxin-3 in HEK293T cells. Cells were processed as described under ‘Materials and Methods’ for visualization under confocal microscope. Arrows indicate the recruitment of PJA1 to ataxin-3 aggregates. (Di) PJA1 colocalizes with 83QHTT aggregate. PJA1 cloned in pEGFP-C1 vector was transiently transfected with pDsRed-83QHTT in HEK293T cells. At 24 hours, cells were processed for confocal microscopy. Colocalization of PJA1 with HTT aggregates was indicated. (Dii) pEGFP-C1/ pDsRed empty vector (EV) and PJA1 alone showed diffused localization in HEK293T cells.

Predictive interactions between PJA1 and ataxin-3 were also indicated by in-silico studies using STRING and BIOGRID databases. To ascertain the interaction between PJA1 ubiquitin ligase and ataxin-3 proteins co-immunoprecipitation was next performed with GFP-ataxin-3 and HA-PJA1. To rule out the nonspecific interaction with GFP and PJA1 initial control experiment was carried out by immunoprecipitating GFP and then probing for PJA1 or vice versa in HEK293T (Human embryonic kidney) cells. No interaction was observed between GFP protein expressed from empty vector and PJA1 (Figure 3A). Subsequently, co-immunoprecipitation was carried out by immunoprecipitating PJA1 and probing for ataxin-3 or vice versa in both HEK293T cells (Figure 3, B and C) and N2A cells (Figure 3, D and E). Robust interaction with not only normal 20QT ataxin-3 protein, but also with 80QT or full length 130Q protein was observed with PJA1. Moreover, to further confirm specificity of interaction between PJA1 and ataxin-3, we performed co-immunoprecipitation experiments between overexpressed PJA1, RING domain deleted PJA1 and endogenous ataxin-3 in HEK293T cells by immunoprecipitating ataxin-3 (Figure 3G) or PJA1 (Figure 3F). A weak interaction between endogenous ataxin-3 with PJA1, but not with its RING deletion counterpart was observed, indicating specificity of association of PJA1 and ataxin-3. Experiments checking the interaction between PJA1 with other polyQ protein huntingtin also produced similar result. PJA1 co-immunoprecipitated with normal (16Q HTT) and polyQ expanded (83Q HTT) huntingtin proteins (Figure 3, I and H); and not with DsRed protein expressed from empty vector (Figure 3H).

**FIGURE 3.**
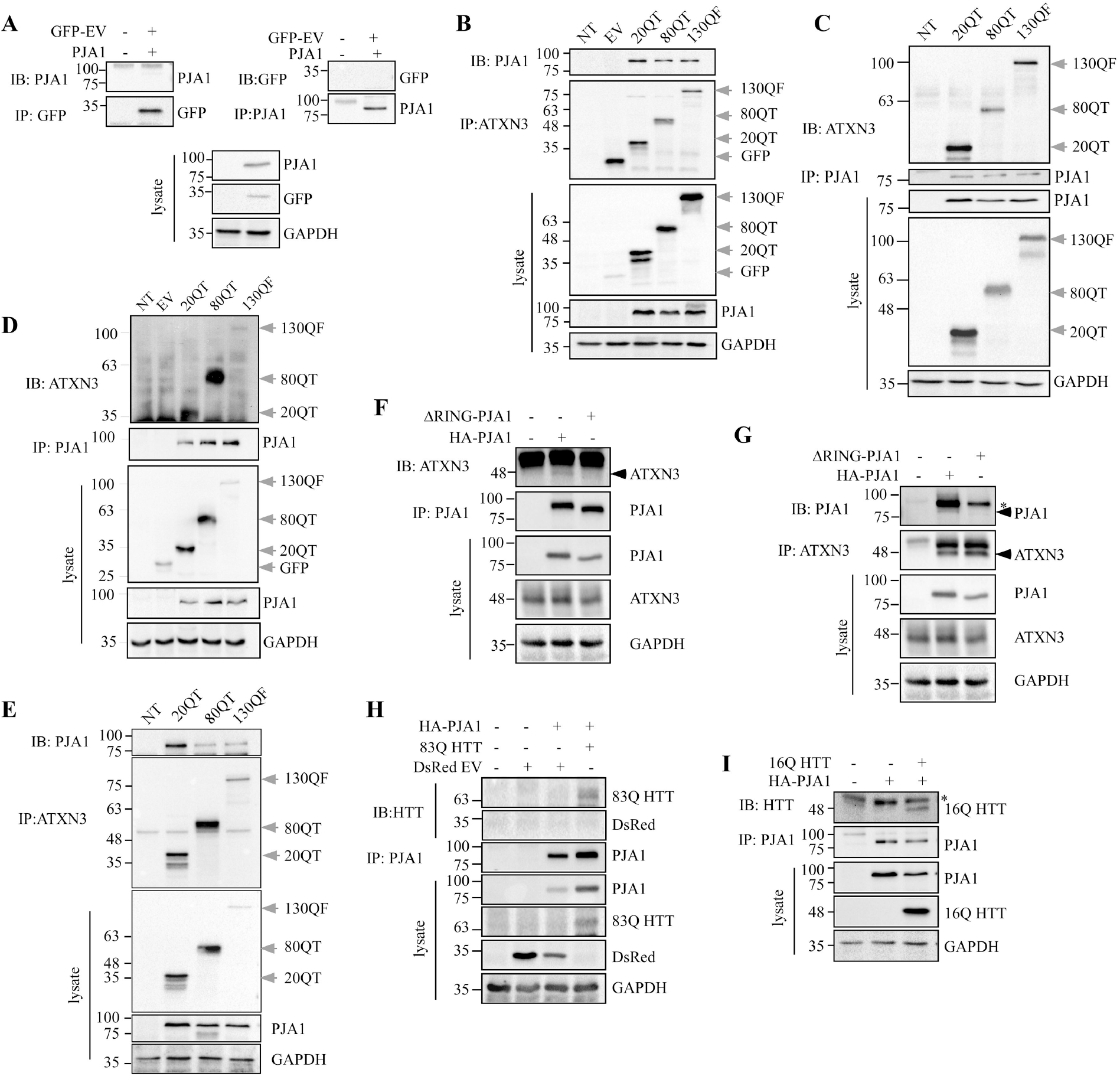
PJA1 interacts with normal and polyQ expanded proteins. (A-E) PJA1 interacts with ataxin-3 proteins. (A) PJA1 does not interact with GFP protein. GFP empty vector (EV) and HA-PJA1 constructs were co-transfected in HEK293T cells. Cell lysate was immunoprecipitated with either anti-GFP or anti-HA antibody followed by western blotting and probing with anti-HA and anti-GFP antibody respectively. (B-D) 20Q truncated (20QT), 80Q truncated (80QT) and 130Q full length (130QF) ataxin-3 in pEGFP-N1 vector were co-transfected with HA-tagged PJA1 (pCMV-HA-PJA1) in HEK293T cells (A and B) and Neuro2A cells (C and D) respectively. 24 hours after transfection, cells were lysed and subjected to immunoprecipitation either with anti-HA antibody (A and C) or with anti-GFP antibody (B and D). The blots were consecutively probed with anti-GFP and anti-HA antibody for detection of ataxin-3 and PJA1 respectively. The total lysate shows the expression of the respective proteins. (F and G) PJA1 interacts with endogenous ataxin-3 protein. HA-tagged PJA1 and ΔRING-PJA1 were transfected in HEK293T cells. 24 hours after transfection, cells were lysed and subjected to immunoprecipitation using anti-SCA3 (F) or anti-HA antibody (G). The blots were consecutively probed with respective antibodies. The total lysate shows the expression of the respective proteins. (H and I) PJA1 interacts with huntingtin protein. N-terminal fragment of 16Q huntingtin (16QHTT) protein in pEGFP-C1 vector (H) and 83QHTT in pDsRed vector (I) were co-transfected with HA-PJA1 in HEK293T cells. Cells were harvested and processed for immunoprecipitation with anti-HA antibody. The blots were successively probed with anti-GFP/anti-DsRed and anti-HA antibody. * indicates non-specific protein. GAPDH is the loading control for all the experiments. Results are representative of three independent experiments. NT and EV represent non-transfected cells and empty vector respectively.

### PJA1 suppresses formation of polyglutamine expanded aggregates

PJA1 is a Ring-H2 finger ubiquitin ligase abundantly expressed in the brain (Yu *et al.*, 2002). With its RING finger motif it promotes degradation of MAGED1 and EZH2 proteins by the ubiquitin proteasome system (Sasaki *et al.*, 2002; Consalvi *et al.*, 2017). PJA1 induces learning in the basolateral amygdala during formation of fear memory which is in coherence to the study establishing the role of PJA1 in cognitive disorders like X-linked mental retardation (Stork *et al.*, 2001; Yu *et al.*, 2002). One of the prominent clinical manifestations of polyQ disorders is cognitive defect (Gardiner *et al.*, 2019), which could be due to dysregulation of PJA1 ultimately leading to toxic accumulation of misfolded polyQ proteins. To examine whether PJA1 can mobilize polyQ aggregates we checked, the formation of ataxin-3 (80QT/130QF) aggregates upon ectopic expression of PJA1 in HEK293T cells by fluorescence microscopy. We noticed a 60-70% reduction in the number of 80QTand 130QF (Figure 4Ai) aggregates counted per 100 GFP positive cells co-expressing ataxin-3 and PJA1. Similar result was obtained for 83QHTT when PJA1 was over-expressed (Figure 4C). Notably, reduced overall fluorescence intensity of 16QHTT was observed in PJA1 expressing condition. This data was further supported by decreased levels of ataxin-3 (Figure 4Aii) and huntingtin proteins as seen in the western blot analysis (Figure 4Ciii). The above results suggest a probable role of PJA1 in mediating degradation of polyQ proteins by utilizing its ubiquitin ligase activity. To confirm that possibility we deleted RING finger motif of PJA1 or mutated its conserved His residue (H553) to Ser and checked its capacity in reducing ataxin-3 aggregates. Our result showed that RING-deleted PJA1 was unable to reduce 80QT ataxin-3 aggregates and its protein level in comparison to WT PJA1, whereas, RING mutant (H553S) PJA1 worked to a lesser extent (Figure 4B). This observation suggests that reduction of polyQ aggregates is linked with PJA1 ubiquitin ligase activity mediated by its RING finger motif.

**FIGURE 4.**
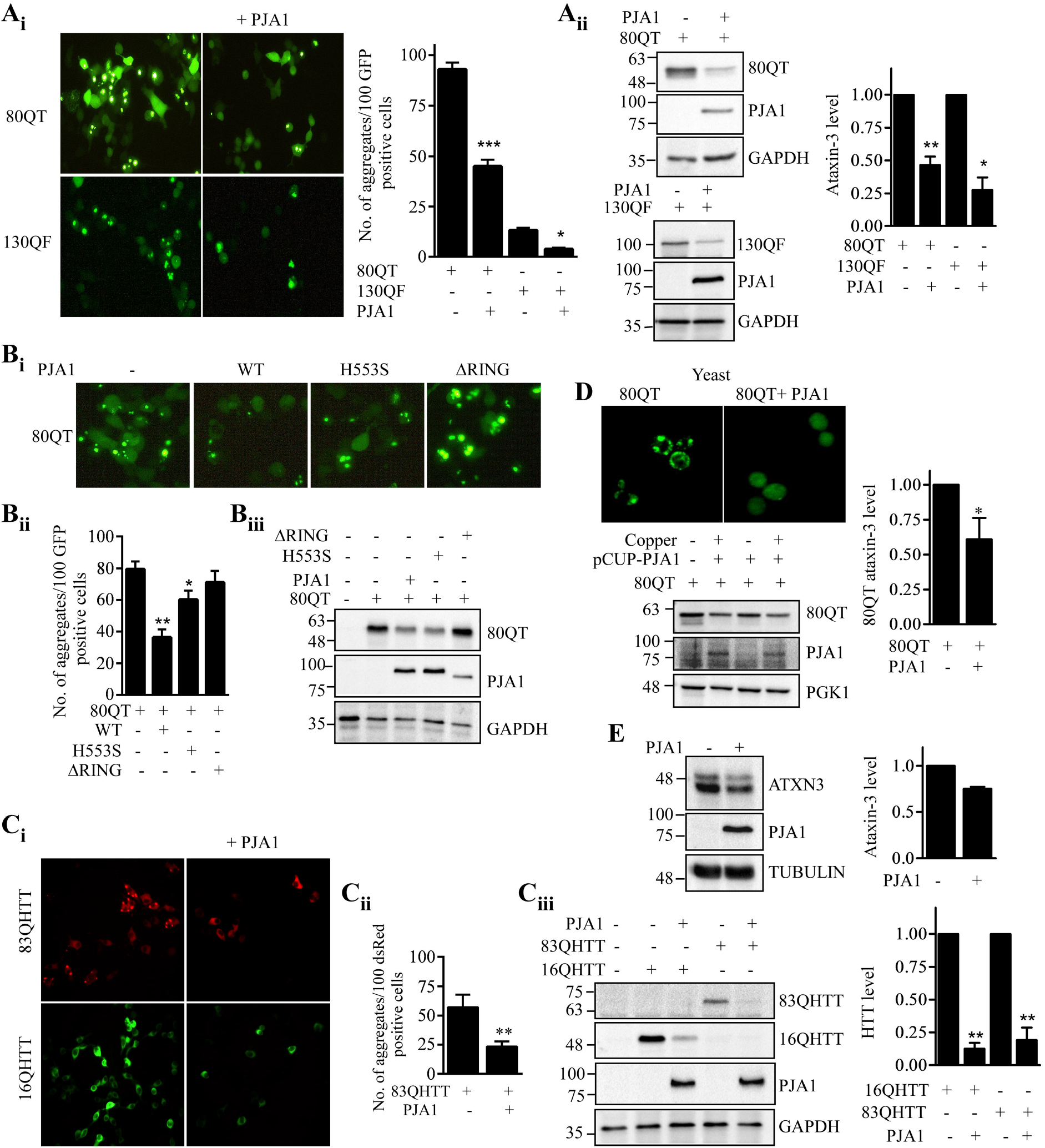
PJA1 suppresses formation of polyglutamine expanded aggregates. (A) PJA1 reduces ataxin-3 aggregates. Ai. HA-tagged PJA1 and the corresponding empty vector were transfected with 80QT and130QF ataxin-3 into HEK293T cells. Transfected cells were viewed under fluorescence microscopy. Aggregates were counted per 100 GFP positive cells from 10 fields randomly chosen and plotted in a bar graph. Values are the mean ± S.D. Significance was calculated by t-test, p<0.05 (*), p< 0.001 (***). (Aii) The lysates from (Ai) were subjected to immunoblotting with anti-GFP and anti-HA antibody for detection of ataxin-3 and PJA1 respectively. The bands were quantified, normalized with GAPDH and plotted as bar diagram. Error bar represents SD of three independent experiments. Significance was calculated by t-test, *, ** indicate p<0.05 and p<0.01 respectively. (B) Impaired PJA1 is unable to suppress aggregation of ataxin-3. 80QT ataxin-3 was transfected alone or in combination with WT, H553S and ΔRING-PJA1 in HEK293T cells. Formation of ataxin-3 aggregates was viewed under a fluorescence microscope (Bi). The aggregates were counted as described earlier and the quantification was plotted in a bar graph (Bii). *, ** indicate p<0.05 and p<0.01 respectively. The same cells were subjected to immunoblotting with anti-GFP, anti-HA and anti-GAPDH antibodies for expression of the corresponding proteins (Biii). (C) PJA1 reduces aggregation of HTT protein. HA-PJA1 was transfected with pDsRed-83QHTT/pEGFP-C1-16QHTT in HEK293T cells (Ci). The 83QHTT aggregates were quantified and plotted as bar diagram (Cii). Error bar represents SD, p<0.01. Western blot analysis shows the expression of the respective protein in cell lysates with anti-GFP, anti-DsRed, and anti-HA antibodies (Ciii). GAPDH serves as loading control. The bands were quantified, normalized with GAPDH and plotted as bar diagram. Error bar represents SD of three independent experiments. Significance was calculated by t-test, p<0.01. (D) PJA1 reduces ataxin-3 aggregation in yeast system. 80QT ataxin-3 and PJA1 were cloned into yeast inducible vectors under galactose and copper promoters respectively. PJA1 expression was promoted by 300μM copper for 2 hours prior to induction of ataxin-3 with 2% galactose for 6 hours. The cells were then subjected to fluorescence microscopy. The same cells were processed for immunoblotting with anti-GFP and anti-HA for detection of ataxin-3 and PJA1. PGK1 indicates loading control. The bands were then quantified, normalized with PGK1 and plotted as bar diagram. Error bar represents SD of three independent experiments. Significance was calculated by t-test, p<0.05. (E) PJA1 reduces endogenous ataxin-3 levels. PJA1 was transfected into HEK293 cells. Cell lysates were subjected to western blotting and endogenous ataxin-3 was detected with anti-spinocerebellar ataxia type 3 1H9 antibody. Tubulin was used as the loading control. The bar diagram was plotted from band intensities calculated and normalized with loading control from two independent experiments.

To further confirm PJA1 driven reduction of polyQ aggregates we used baker’s yeast *Saccharomyces cerevissiae* which is devoid of both the proteins. Being a eukaryote, *Saccharomyces cerevissiae* serves as an effective model and is widely used for many protein misfolding diseases including neurodegenerative diseases (Miller-Fleming *et al.*, 2008). We cloned ataxin-3 and PJA1 in yeast under galactose and copper inducible promoters respectively. PJA1 expression was promoted by the addition of 300μM copper prior to ataxin-3 expression with 2% galactose. The status of the ataxin-3 aggregates was then checked by fluorescence microscopy. We noticed an overall decrease in the number of ataxin-3 aggregates in yeast cells expressing PJA1 which was further maintained by significant reduction in the protein level as well (Figure 4D). Furthermore, reduction of endogenous level of ataxin-3 protein was observed upon PJA1 expression in HEK293 cells (Figure 4E). On contrary, when endogenous PJA1 was silenced using shRNA targeted to PJA1 (as shown by RT-PCR due to poor sensitivity in detecting endogenous PJA1 with antibody (Figure 5A)) in N2A cells over-expressing normal and polyQ expanded ataxin-3, approximately five-six-fold increment in protein levels of the respective ataxin-3 proteins was detected (Figure 5B). The knockdown of endogenous PJA1 levels also resulted in significant increase in the number of polyQ expanded ataxin-3/huntingtin aggregates (Figure 5, C and D). Similar results were obtained by using siRNA targeted to PJA1 in N2A cells (Figure 5, E-G). Silencing of PJA1 enhances 20QT and 80QT ataxin-3 protein level as seen by western blot analysis and also by fluorescence microscopy. Altogether, these results suggest that PJA1 modulates polyQ proteins and their aggregates by utilising its ubiquitin ligase activity.

**FIGURE 5.**
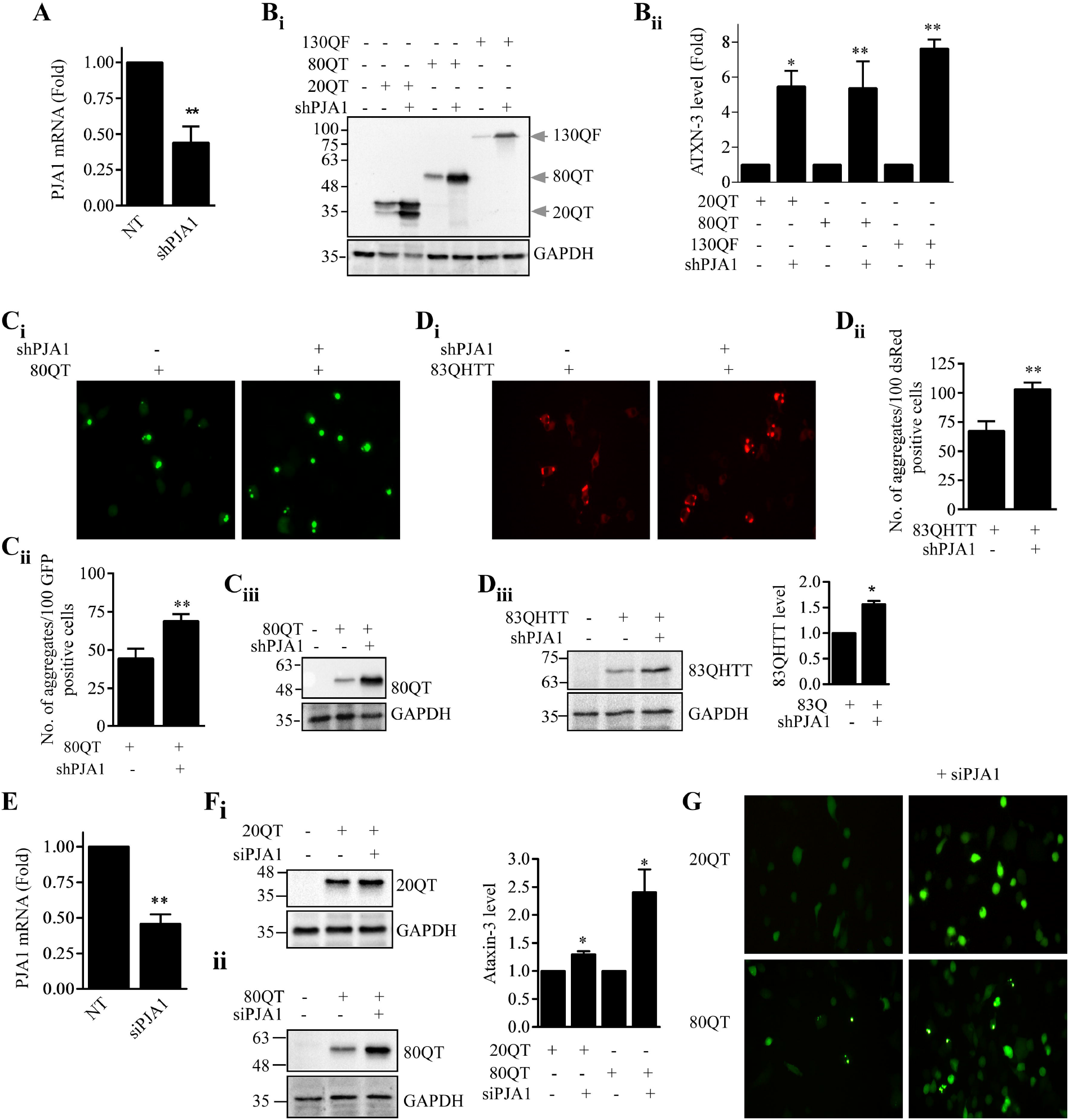
PJA1 silencing in Neuro2A cells enhances formation of polyQ aggregate. (A) shRNA mediated down regulation of PJA1 transcript level. Neuro2A cells were transiently transfected with short hairpin RNA (shRNA) targeted to PJA1. Cells were harvested after 48 hours. Total RNA was isolated and subjected to quantitative real-time PCR. Data were collected from three independent experiments and normalised to the levels of GAPDH. Values are the mean ± SD. Significance was calculated by t-test, p< 0.01. (B) PJA1 down regulation enhances the level of ataxin-3 proteins in Neuro2A cells. The protein level of 20QT, 80QT and 130QF ataxin-3 was assayed in PJA1 silenced condition by western blot analysis. The blots were probed with anti-GFP antibody for detection of ataxin-3. GAPDH is the loading control. The bands were quantified, normalized with GAPDH and plotted as bar diagram. Error bar represents SD of three independent experiments. Significance was calculated by t-test, *, ** indicate p<0.05 and p<0.01 respectively. (C and D) Downregulation of PJA1 enhances ataxin-3 and HTT aggregates. 80QT ataxin-3 (C) or 83QHTT (D) plasmids were transfected in Neuro2A cells upon silencing of PJA1 with shRNA. After 48 hours cells were viewed under fluorescence microscopy. The aggregates were counted and plotted in a bar graph. Values are the mean ± SD, Significance was calculated by t-test, p<0.01. Same cells were processed for immunoblotting and the blots were probed with anti-GFP, anti-DsRed and anti-GAPDH antibodies. The 83QHTT bands were similarly quantified and plotted as bar diagram, the error bar representing SD of three independent experiments. Significance was calculated by t-test, p<0.05. (E) siRNA mediated down regulation of PJA1 transcript level. Neuro2A cells were transiently transfected with scramble or siRNA targeted to PJA1. Cells were harvested after 48 hours. Total RNA was isolated and subjected to quantitative real-time PCR. Data were collected from three independent experiments and normalised to the levels of GAPDH. Values are the mean ± SD. Significance was calculated by t-test, p< 0.01. (F) siRNA mediated down-regulation of PJA1enhances ataxin-3 protein level in Neuro2A cells. The protein level of 20QT (Fi), 80QT (Fii) ataxin-3 was assayed in PJA1 silenced condition by western blot analysis. The blots were probed with anti-GFP antibody for detection of ataxin-3. GAPDH is the loading control. The bands were quantified and normalized with GAPDH and plotted as a bar diagram. Error bars represent a cumulative of three experiments performed independently. Significance was calculated by t-test, p<0.05. (G) The same cells were viewed under fluorescence microscope, where enhanced overall fluorescence and aggregation in PJA1 silenced condition was observed for 20QT and 80QT ataxin-3 respectively.

### PJA1 facilitates degradation of wild type and polyglutamine expanded ataxin-3 by promoting ubiquitination

Based on previous reports of substrate degradation by PJA1 ubiquitin ligase (Sasaki *et al.*, 2002; Consalvi *et al.*, 2017), and diminution of steady state expression of ataxin-3 and huntingtin proteins in mammalian and yeast cells in presence of PJA1 (Figure 4, A, C and D), we wanted to investigate whether PJA1 aided in the degradation of polyQ proteins. Moreover, unaltered endogenous ataxin-3 mRNA level in PJA1 overexpressed cells ruled out possible transcriptional down-regulation of ataxin-3 by PJA1 (Figure S1) which further support PJA1’s function in reducing ataxin-3 at the post-translational level. Tracking the turnover of polyQ proteins in mammalian cells was not pursued as PJA1 is a short lived protein promoting self ubiquitination (Zoabi *et al.*, 2011). Instead, substitute experiments in mammalian cells (N2A) were performed wherein PJA1 was overexpressed in ascending concentrations and ataxin-3 was found to be reduced in a concentration dependent manner (Figure 6, A and B). Furthermore temporal expression of PJA1 showed a gradual decrease in ataxin-3 protein levels over time (Figure 6C). These results suggest a correlation between PJA1 over-expression and reduction of ataxin-3 level. To confirm this we pursued galactose shut-off chase of 80QT ataxin-3 in yeast system upon expression of PJA1 from an inducible copper promoter. Enhanced turn-over of ataxin-3 was observed till 4 hr of chase when adequate amount of PJA1 is present that suggests PJA1 mediated degradation of ataxin-3 (Figure 6D). Furthermore, stabilization of 80QT ataxin-3 was observed in cycloheximide chase upon silencing of PJA1 in N2A cells (Figure 6E).

**FIGURE 6.**
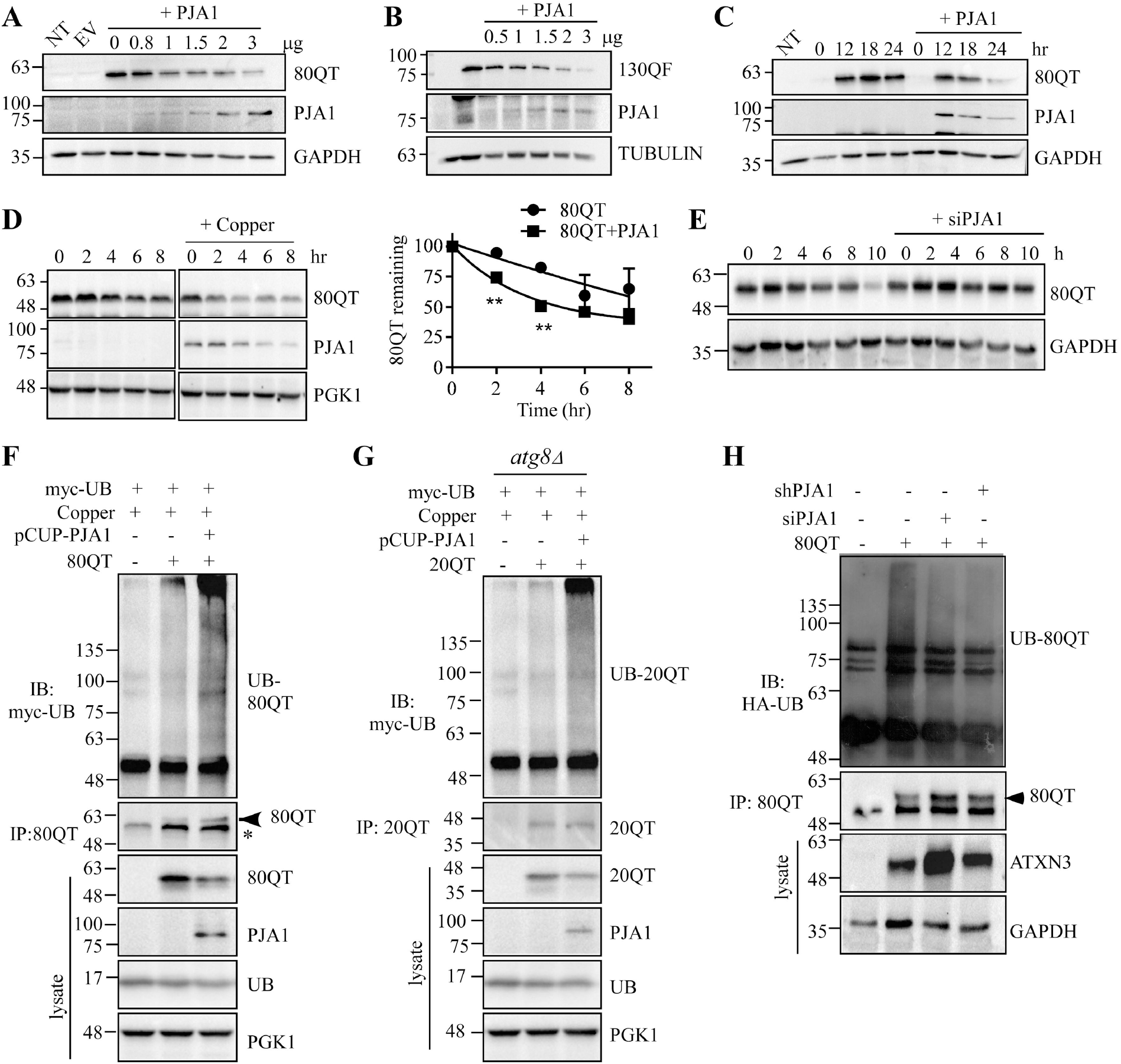
PJA1 promotes degradation of ataxin-3 by promoting its ubiquitination. (A-C) Time and dose dependent expression of PJA1 enhances degradation of ataxin-3 protein. Neuro2A cells were transiently transfected with 80QT (A), and 130QF (B) with varying concentrations of PJA1 as indicated. 24 hours after transfection, ataxin-3 level was detected in cell lysates by western blotting with anti-GFP antibody. C. 80QT ataxin-3 and PJA1 (3 μg) were co-transfected in Neuro2A cells and the cells were collected at varying time as indicated. Western blot analysis shows ataxin-3 level. GAPDH and Tubulin was used as loading controls. (D and E) PJA1 facilitates degradation of 80QT ataxin-3 in mammalian and yeast system. (D) GAL-shut off chase was performed to measure the turn-over of ataxin-3. PJA1 expression was carried out with addition of copper followed by induction of ataxin-3 with 2% galactose for 2 hours. The expression of ataxin-3 was shut off using 2% glucose, and chased for indicated times. Ataxin-3 level was detected by western blotting with anti-GFP antibody. PGK1 is loading control. Data were collected from three separate experiments and normalised to the levels of PGK1. The band intensities were quantified using imageJ software and plotted in the adjacent graph. Error bar indicates SD. (E) Cycloheximide chase of 80QT ataxin-3 was done upon silencing of PJA1 with smartpool siRNA (Dharmacon) targeted to PJA1 in Neuro2A cells as described in method section. 10μg/ml of cycloheximide was used to stop translation and chased for 10 hours. Ataxin-3 level was detected by western blotting with anti-GFP antibody. GAPDH acts as the loading control. (F-H) PJA1 promotes ubiquitination of ataxin-3. (F and G) 80QT and 20QT ataxin-3 plasmids were transformed along with PJA1 and myc-ubiquitin into WT (F) and *atg8Δ* (G) yeast cell respectively. Expression of PJA1 was induced with 300μM copper for 2 hours prior to ataxin-3 induction with 2% galactose for 4 hours. Yeast cells were then harvested and immunoprecipitation was performed using anti-GFP antibody. The blots were probed with anti-myc and anti-GFP antibody for detection of ubiquitinated ataxin-3 and the amount of immunoprecipitated ataxin-3 respectively. Lysate shows the expression of the proteins. PGK1 was the loading control. (H) Silencing of PJA1 reduces ubiquitination of 80QT ataxin-3. PJA1 was silenced in Neuro2A cells either with shRNA or siRNA targeted to PJA1. GFP-80QT ataxin-3 and HA-ubiquitin plasmids were co-transfected and the ubiquitination of 80QT ataxin-3 was detected upon immunoprecipitating ataxin-3 with anti-GFP antibody followed by western blotting with anti-HA antibody. Lysates show the expression of respective proteins. GAPDH is the loading control.

Since degradation is accompanied by tagging substrate molecules with ubiquitin moieties by ubiquitin ligase, we next assessed whether PJA1 was capable of ubiquitinylating ataxin-3 in yeast system. We noticed an increased ubiquitination of 80QT ataxin-3 when PJA1 was expressed in yeast cells from a copper inducible promoter (Figure 6F). Similarly, ubiquitination of 20QT ataxin-3 was drastically increased in *atg8* knockout yeast cells expressing PJA1 (Figure 6G). On contrary, silencing of PJA1 by shRNA or siRNA (validated by RT-PCR) targeted to PJA1 reduces polyubiquitination of 80QT ataxin-3 in N2A cells (Figure 6H). Together, these results corroborate the fact that PJA1 is an E3 ubiquitin ligase which ubiquitinates ataxin-3 and hence promotes its degradation.

Ubiquitin ligase mediated clearance of protein aggregates is powered preferentially by the autophagy pathway (Nixon, 2013). Previously PJA1 function has been shown in degradation of MAGED1 and EZH2 proteins by the UPS (Sasaki *et al.*, 2002; Consalvi *et al.*, 2017). Aligning this result, we also found rescue of EZH2 protein upon inhibition of proteasome by lactacystin in PJA1 expressing HEK293T cells (Figure 7A). Here, we wanted to check whether PJA1 shuttles ubiquitinated polyQ proteins via UPS or autophagy. Interestingly, PJA1 mediated reduction of ataxin-3 protein was rescued when the proteasome or autophagy pathway is blocked (Figure 7, B and C). This result suggests proteasomal as well as autophagic clearance of ataxin-3 facilitated by PJA1. To further validate autophagy mediated degradation of ataxin-3 by PJA1, yeast knockout strains *atg8Δ* was used. 20QT ataxin-3 levels were refurbished in *atg8Δ* strains, confirming the role of PJA1 in promoting degradation of ataxin-3 also through the autophagy pathway (Figure 7D).

**FIGURE 7.**
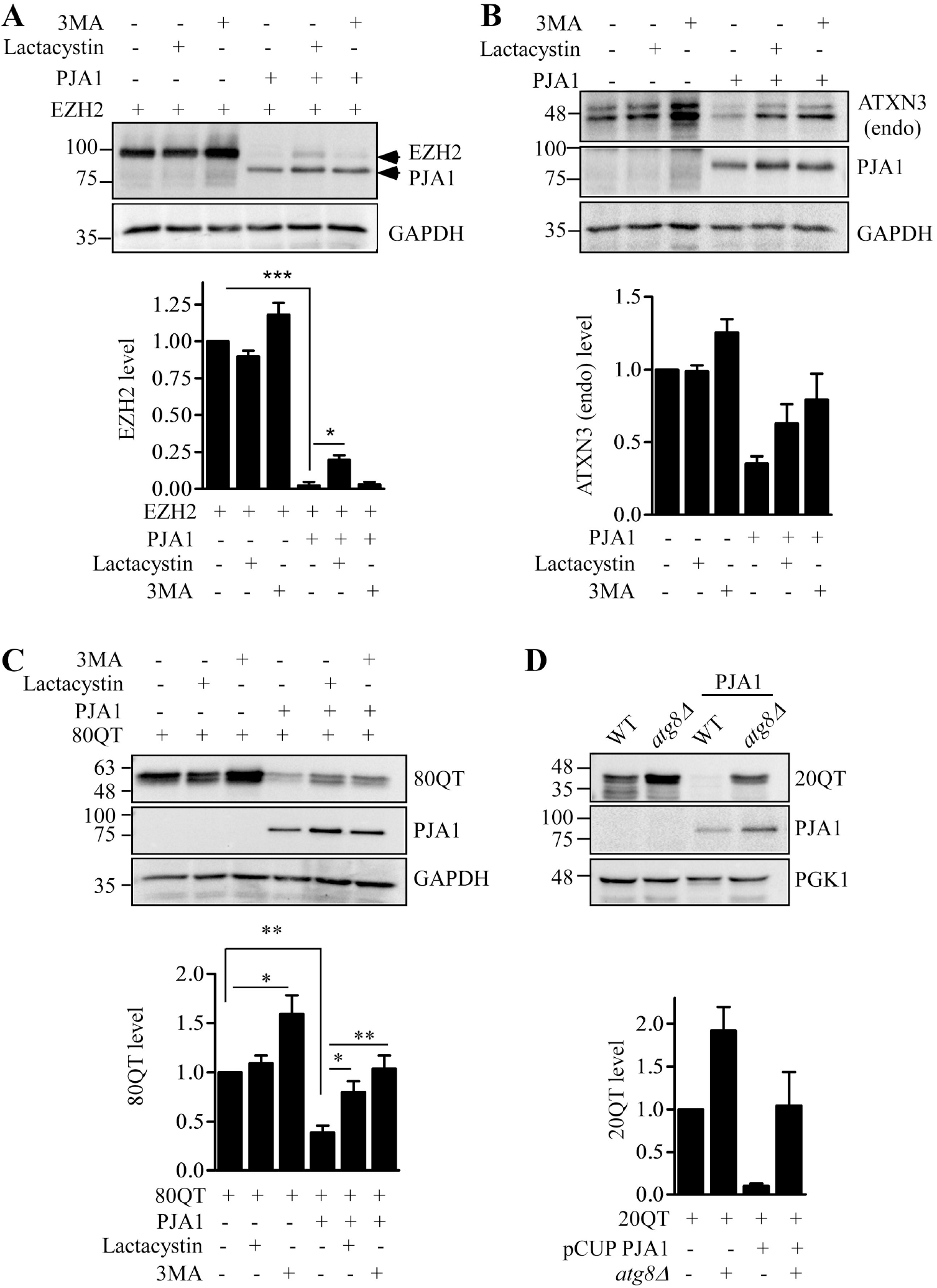
PJA1 facilitates degradation of ataxin-3 through proteasome and autophagy. (A) PJA1 mediated proteasomal degradation of EZH2 protein. EZH2 and PJA1 was co-transfected in HEK293T cells. Cells were treated with either lactacystin (2μM) or 3MA (4nM) for 6 hours before harvesting the cells. EZH2 and PJA1 levels were detected by western blotting with anti-HA antibody. GAPDH is loading control. Data collected from three separate experiments were normalised to the levels of GAPDH. The band intensities were quantified using ImageJ software and plotted in the graph. Error bar indicates SD. Significance was calculated by t-test, p<0.05 (*), p<0.001(***). (B-D) PJA1 clears ataxin-3 via proteasome and autophagy. (B) Endogenous ataxin-3 level was monitored upon overexpression of PJA1 in HEK293T cells treating with lactacystin or 3MA. The graph has been plotted using data from two different experiments. (C) 80QT ataxin-3 and PJA1 was coexpressed in HEK293T. Ataxin-3 level was assayed by western blot analysis upon treating the cells with lactacystin or 3MA. The bands were quantified and normalized with GAPDH and plotted in the graph. Values are the mean ± SD. Significance was calculated by t-test, p<0.05 (*), p<0.01 (**). (D) Knock out of *ATG8* rescued PJA1 mediated degradation of 20QT ataxin-3 in yeast. p315GAL-GFP 20QT ataxin-3 and p426CUP-HA-PJA1 plasmids were transformed into *atg8Δ* yeast strains. PJA1 expression was induced with 300μM copper followed by ataxin-3 with 2% galactose. Over-expressed and endogenous ataxin-3 levels were monitored by western blotting using anti-GFP and anti-SCA3-1H9 antibodies respectively. Data from two separate experiments were plotted as bar diagram. GAPDH (panel A-C) and PGK1 (panel D) were used as loading controls.

### PJA1 suppresses polyglutamine toxicity in yeast and Drosophila model of SCA3

Accumulation of expanded polyQ proteins causes neuronal vulnerability, although the principal reason responsible for its toxicity is currently unknown (Paulson *et al.*, 2017). Evidence suggests polyQ expansion driven acceleration of neuronal aging with altered nuclear integrity and nucleocytoplasmic transport in Huntington’s disease (Gasset-Rosa *et al.*, 2017). Since PJA1 promoted degradation of ataxin-3 protein (Figures 5 and 6), thus it might lead to decreased proteotoxicity associated with accumulation of mutant ataxin-3. To test that we have initially chosen yeast as a model to study polyglutamine mediated toxicity. Expression of 80QT ataxin-3 from galactose inducible promoter in yeast causes growth retardation of yeast cells as evident from a spot assay. However, expression of PJA1 in these cells rescues yeast toxicity suggesting suppression of polyQ pathogenesis by PJA1 (Figure 8A). We further validated this result in *Drosophila* model of SCA3. The transgenic models used in this study take advantage of the GAL4-UAS system, in which cDNAs encoding human mutant ataxin-3 (truncated 78Q, Bloomington ID- 8150) are under control of the heterologous yeast Upstream Activating Sequence (UAS) promoter element which respond to driver lines expressing the GAL4 transcription factor. 78QT ataxin-3 under the GMR-GAL4 system shows severe degenerative external eye phenotype, however, GMR-GAL4-PJA1 transgenic fly was embryonically lethal. Thus, Rhodopsin1-GAL4 (Rh1) line was selected that is expressed following photoreceptor maturation and is active only after completion of eye development beginning in late pupal stages. Transgenic PJA1 fly was stable under GAL4-Rh1 system and PJA1 expression levels under GAL4-Rh1 were checked (Figure S2). Cryo-sectioning of transgenic *Drosophila* eyes followed by hematoxylin/eosin staining was performed. In young Rh1-78QT (8-10 day) or Rh1-PJA1, retinal sections did not exhibit specific defects in ataxin-3 expressing flies (Figure 8B). However, in older Rh1-78QT flies (35-40 day), the retinas exhibited striking degenerative phenotypes with disorganized ommatidia and tissue dissociation associated with the appearance of vacuoles as mentioned in previous studies (Sowa *et al.*, 2018). The Rh1-78QT flies with over-expressed PJA1 had however far less vacuole formation when compared to the Rh1-78QT, establishing PJA1’s role in mitigating toxicity caused by polyQ expanded ataxin-3 (Figure 8C). We additionally performed immunofluorescence of these retinal cryosections to confirm expression of ataxin-3 and PJA1. We obtained similar morphological differences as observed in Fig. 8C. Furthermore, the aggregates formed in case of Rh1-78QT flies were considerably lesser in flies overexpressing PJA1 which reconfirms PJA1’s role in degradation of polyQ proteins (Figure 8, Diii-v).

**FIGURE 8.**
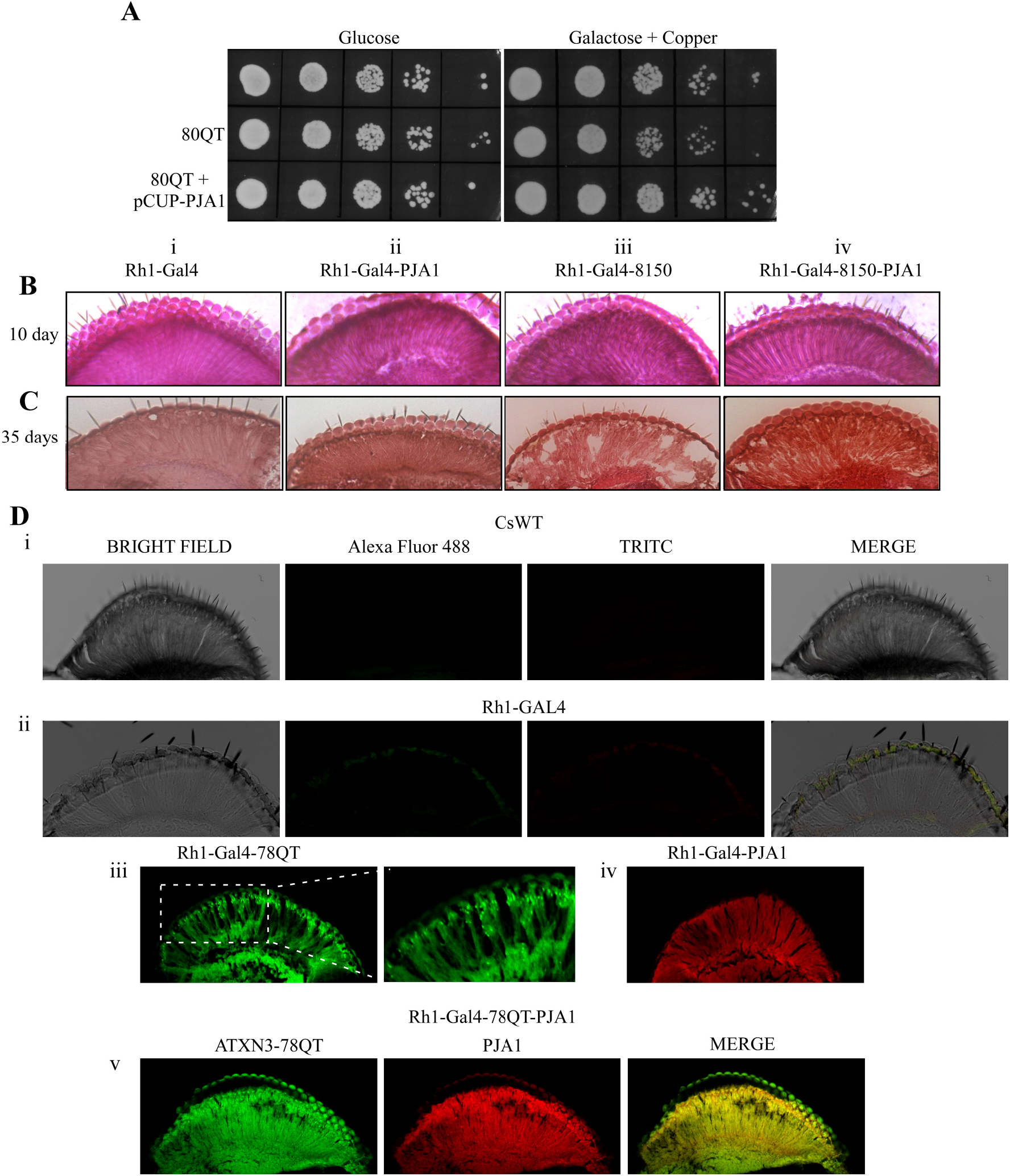
PJA1 suppresses polyglutamine toxicity in yeast and *Drosophila* model of SCA3. (A) PJA1 suppresses polyQ mediated toxicity in yeast. Yeast cells harbouring p315GAL 80QT ataxin-3 and p426CUP-PJA1 were grown overnight in 2% raffinose. PJA1 was induced with 300μM copper for 2 hours followed by induction of ataxin-3 with 2% galactose. The cultures were serially diluted and spotted on to either YPD or YP with 2% galactose and 100μM copper. The plates were incubated at 30°C. Growth of yeast cells were observed after 48 hours. (B and C) PJA1 rescues polyQ induced toxicity in *Drosophila* eye. (Bi-iv) Retinal sections of *Drosophila* eye of 8-10 day old fly. Retinal sections of mutant ataxin-3 expressing transgenic *Drosophila* eye showed identical phenotype with the control fly. Notably, *Drosophila* expressing PJA1 did not exhibit any change in retinal morphology. (C) 35-40 day old fly developed retinal vacuole formation and photoreceptor disorganization induced by mutant ataxin-3 (Ciii). Over-expression of PJA1 in mutant ataxin-3 expressing *Drosophila* results in retinal sections with lesser vacuoles and an overall rescue of phenotype. (n=10 per genotype). (D) Immunofluorescence of the retinal cryosections of the 35-40 day old flies expressing mutant ataxin-3 (D_iii_), PJA1 (D_iv_) alone and coexpression of mutant ataxin-3 and PJA1 (D_v_). The CsWT (D_i_) and Rh1-GAL4 (D_ii_) flies were taken as control.

## DISCUSSION

Eukaryotic cells are equipped with protein quality control (PQC) machinery which eliminate damaged or misfolded proteins and relieve cells from proteotoxic stress (McKinnon and Tabrizi, 2014). However when misfolded proteins overwhelm the capacity of the clearance machinery, they accumulate as intracellular aggregates which are toxic for the cell. Polyglutamine (polyQ) repeat proteins undertake abnormal conformation when its polyQ tract exceeds beyond the threshold level and form intracellular inclusions/aggregates, which account for cellular toxicity, characterised by microtubule destabilization, cytoskeleton collapse, transcriptional dysfunction and finally cell death (Trushina *et al.*, 2003). The components of the PQC, the molecular chaperones try to refold them, but on failure, shuttle them to E3 ubiquitin ligases to be tagged by ubiquitin and eventually degraded by the ubiquitin proteasome pathway or autophagy. There are several E3 ligases viz. CHIP, Parkin, Gp78, E6-AP, MGRN1, ITCH and Malin, that aid in the clearance of polyglutamine proteins and their aggregates hence mitigating cytotoxicity (Tsai *et al.*, 2003; Jana *et al.*, 2005; Garyali *et al.*, 2009; Ying *et al.*, 2009; Chhangani and Mishra, 2013; Mishra *et al.*, 2013; Chhangani *et al.*, 2014). In the present study, we described the function of Praja1 (PJA1), a RING-H2 ubiquitin ligase highly expressed in brain, in degradation of polyQ proteins. PJA1 associates with polyQ proteins and leads to reduction of polyQ aggregates and cytotoxicity in cellular and *Drosophila* models of SCA3.

Proteostasis collapse or proteostatis network (PN) disruptions are fundamental in several age-onset neurodegenerative disorders (Hipp *et al.*, 2014). However, the reason behind this collapse, and how it contributes to disease onset or progression, is still elusive. One hypothesis is that chronic expression of aggregation–prone polyQ expanded proteins, α-synuclein or Aβ leads to proteasomal and autophagy impairment which drive the cellular disruptions observed in most neurodegenerative disorders such as HD, PD and AD (Gregori *et al.*, 1995; Keller *et al.*, 2000; McNaught and Jenner, 2001; Komatsu *et al.*, 2006; Winslow *et al.*, 2010; Hipp *et al.*, 2012). Our observation strengthens the hypothesis. We found reduced levels of PJA1 transcript and protein in cells expressing polyQ proteins. The simple explanation could be the fact that polyQ aggregates sequester transcription factors such as nuclear factor kappa-light-chain-enhancer of activated B cells (NF-κB), CREB binding protein (CBP), TATA box binding protein (TBP), which as a consequence lose their normal cellular functions (Shimohata *et al.*, 2000; Dunah *et al.*, 2002; Schaffar *et al.*, 2004; Goswami *et al.*, 2006). The aberrant function of transcription factors thus contributes to enhanced pathogenesis of polyQ diseases. These findings also led us to speculate the interaction of PJA1 with polyQ proteins as polyQ aggregates voraciously sequester components of the PQC: ubiquitin ligases such as MGRN1, CHIP and Malin, and hence unavailability of functioning cellular surveillance system may also be responsible for disease pathogenicity. Likewise, we find PJA1 interaction with wild type as well as mutant ataxin-3/huntingtin proteins and also their aggregates (Figures 2 and 3). A recent report establishing PJA1’s role in reducing TDP43 aggregate formation and pathogenesis associated with ALS and FTLD also supported our hypothesis (Watabe *et al.*, 2020).

These results impelled us to identify PJA1’s functional role in the proteostasis of polyQ proteins. Our observations established a marked decrease in the number of polyQ aggregates by PJA1, which was not detected in case of RING domain deleted PJA1 (Figure 4). The finding was confirmed by shRNA/siRNA mediated knock-down of PJA1, which resulted in increase in the polyQ aggregate count and their protein levels (Figures 5 and 6G). This lead us to postulate PJA1’s role as an E3 ubiquitin ligase in degradation of ataxin-3 and hence the removal of aggregates. Previously, PJA1’s role had been established in degradation of proteins such as MAGE-D1/Dlxin-1, EZH2 involved in osteoblast differentiation and myogenesis respectively (Sasaki *et al.*, 2002; Consalvi *et al.*, 2017). Over-expression of PJA1 has been found in cancers such as glioblastomas and gastrointestinal cancer where adaptor protein ELF of tumour suppressor SMAD3 is degraded (Saha *et al.*, 2006; Chen *et al.*, 2020). But function of PJA1 in degradation of polyQ proteins had not been studied previously. Ubiquitin ligases work by tagging ubiquitin to substrates for their recognition by the proteosomal machinery/autophagy for their degradation (Deshaies and Joazeiro, 2009). In line with that, we found enhanced ubiquitination of ataxin-3 in presence of PJA1, whereas diminished ubiquitinated ataxin-3 when PJA1 was silenced, indicating PJA1’s ability to ubiquitinate polyQ proteins (Figure 6). Notably, PJA1 mediated degradation of polyQ proteins occurs via proteasome as well as lysosomal pathway although further study is needed to establish the preferential pathway or dependency on substrates.

Clearance of misfolded proteins from cell leads to reduction in its accumulation and associated cellular toxicity. Our results thus prompted us to examine the consequential effects of removal of aggregates and degradation of polyQ proteins by PJA1. Previously CHIP, UBR5, UBE3A have been shown to mitigate polyQ induced toxicity in cellular as well as animal models of polyQ diseases (Miller *et al.*, 2005; Maheshwari *et al.*, 2014; Koyuncu *et al.*, 2018). Similar to these ubiquitin ligases, PJA1 is capable in reducing the growth sensitivity of yeast associated with polyQ protein and rescuing the retinal degeneration in *Drosophila* model of SCA3 (Figure 8). This result suggests a cytoprotective response of PJA1 against polyglutamine induced toxicity in yeast as well as *Drosophila*.

Interestingly, studies panning all polyQ disorders hypothesize presence of tissue specific proteostasis network (PN). PN heterogeneity encompasses all classes of chaperone, autophagy mediators, UPS components, and stress response regulators and they exhibit profoundly altered expression patterns between tissues (Powers *et al.*, 2009; Tebbenkamp and Borchelt, 2010; Guisbert *et al.*, 2013). Previous studies have also hinted at how PN heterogeneity can influence disease presentation and progression (Tagawa *et al.*, 2007; Tsvetkov *et al.*, 2013). Hence deciphering the entire complex PN in polyQ disorders might help us understand the major chaperones and degradative components at work. And, since the requirements of the PQC by the substrates vary based on tissue type, there might be a robust PN working in the brain which modulates normal or neurodegenerative disease related proteins. PJA1 serves as a crucial E3 ligase of the brain proteome which is recruited to TDP43 (Watabe *et al.*, 2020) and polyQ proteins like ataxin-3 and huntingtin for their degradation, and hinders their aggregation; and thus dysfunction of this E3 ligase might result in pathogenesis and onset of neurodegeneration. In conclusion, our results suggest that modulation of PJA1 might serve as potential therapy for polyQ disorders and result in weakened pathogenesis of the diseases.

## MATERIALS AND METHODS

### Strains, Reagents and antibodies

Yeast strain, BY4741 (MATa his3Δ0 leu2Δ0 met15Δ0 ura3Δ0) was used in this study. Autophagy impaired *atg8::Kanmx* was obtained from yeast knock out library. Lactacystin, 3-Methyladenine (3MA) were purchased from Sigma. The following primary antibodies were used: mouse monoclonal antibodies against Spinocerebellar ataxia Type 3 SCA3-1H9, DsRed (Merck-Millipore), FLAG, HA (Sigma), GFP (Roche), Ub (Cell Signaling Technology), PGK1 (Invitrogen); rabbit monoclonal antibodies against: α-Tubulin, GAPDH (BioBharati Life Sc, India), MYC, GFP (Cell Signaling Technology), PJA1 (Genetex). The secondary antibodies, goat anti-mouse IgG-HRP antibody and anti-rabbit IgG-HRP were obtained from Jackson Laboratory.

### Plasmid constructs

Ataxin-3 constructs (pEGFP-N1-20QT, pEGFP-N1-80QT and pEGFP-N1-130QF) were kindly provided by Prof. Nihar Ranjan Jana (IIT KGP, India). Mouse Praja1 in vectors pCMV-HA-PJA1, pEGFP-C1-PJA1 were a gift from Dr. Oliver Stork (Otto-von-Guericke University Magdeburg, Germany). Exon1 of pEGFP-C1-16Q and pDsRed-C1-83Q huntingtin were also obtained from Dr. Debashis Mukhopadhyay (Saha Institute of Nuclear Physics, India). The following yeast constructs p315GAL-20QT-GFP, p315GAL-80QT-GFP were generated by standard PCR methods. 20QT and 80QT were first amplified by PCR using primers 5′FLAG BamH1-CGCGGATCCATGGACTACAAAGACGATGACGACAAGCAAGGTAGTTCCAG and 5′ Not1 TTTTCCTTTTGCGGCCGCCTAGATCACTCC (without stop codon). The full ORF (20QT/80QT) was then cloned into p315GAL plasmid digested with BamH1 and Not1. The ORF of GFP was digested from YEp53 plasmid using Not1 restriction enzyme and inserted after 20QT and 80QT. PJA1 was amplified from pEGFP-C1-PJA1 using primers 5′HA HindIII

CCCAAGCTTATGTACCCATACGATGTTCCAGATTACGCTAGCCACCAGGAAAGG A and 5′Xho1 CCGCTCGAGTTAGAGCGGGGGAGGGAAC and cloned into p426CUP vector. The deletion mutants and point mutations of PJA1 were obtained by using standard PCR and site directed mutagenesis protocols. pDsRed-C1-PJA1 was generated by subcloning from pEGFP-C1-PJA1. All constructs were confirmed by sequencing. Plasmids used in this study are shown in Table 1 of supplemental material.

### Cell culture and transfection

Human Embryonic Kidney 293T (HEK293T), HEK293 or Mouse Neuroblastoma (N2A) cells were obtained from National Centre for Cell Science (NCCS), Pune, India. The cells were cultured in Dulbecco’s modified Eagle’s medium (DMEM) (HiMedia Laboratories, India) containing 10% foetal bovine serum (FBS) (GIBCO) in humidified chamber with 5% CO2. The plasmid DNA was transfected using Lipofectamine LTX with plus reagent (ThermoFisher) or PEI (Polyetheleneimine, Polysciences Inc.) at 70% confluence in DMEM without serum. shRNA/siRNA transfection was carried out with Lipofectamine 2000 (ThermoFisher) according to the manufacturer’s protocol.

### Western blot analysis

Cells were first lysed in RIPA buffer containing 50 mM Tris–HCl (pH 7.5), 150 mM NaCl, 1 mM EDTA, 0.1% SDS, and 1 mM PMSF with protease inhibitor cocktail. Cell lysates were cleared at 12,000g for 20 min at 4°C. Proteins were estimated using the BCA protein estimation kit (ThermoFisher) and subsequently separated by 10% SDS-PAGE and transferred onto nitrocellulose membrane (Pall Corporation, NY). The proteins were visualized using ECL detection buffer and image was captured in Chemidoc MP system (BioRad). Image analysis was done by ImageJ software. Statistical analysis was scored using GraphPad Prism software.

### Immunoprecipitation

Cells were lysed in buffer containing 50 mM Tris, pH 7.5, 150 mM NaCl, 0.1% NP-40, 1mM PMSF and protease inhibitor cocktail. Cellular debris was removed by centrifugation at 12,000g for 15 min at 4°C. Protein concentration was estimated by BCA method. The cell lysates were incubated with antibody in IP dilution Buffer (50mM Tris, pH7.5, 150 mM NaCl, 0.1% NP-40, PMSF) overnight at 4°C with gentle rotation. Next day, protein A Sepharose (GE healthcare) bead pre-equilibrated with IP dilution buffer was added and nutated further for 2 hr at 4°C for immunoprecipitation. The beads were then washed three times with IP dilution buffer. Bound proteins were eluted from beads with SDS sample buffer, vortexed, boiled for 5 minutes and analysed by western blotting.

### Microscopic analysis

HEK293T cells were transfected with polyQ proteins ataxin-3 (pEGFP-N1-80QT and 130QF) or huntingtin (16Q and 83Q) alone and with pCMV-HA-PJA1. 24 hours after transfection, cells were viewed under fluorescence microscope at 20X (Leica). Aggregates were counted per 100 GFP positive cells, from 10 randomly chosen fields.

Confocal Imaging-HEK293T cells were grown on cover slips coated with 0.01% poly-L-Lysine. DNA transfection was carried out at 60% confluence. After 24 hours the cells were fixed with 3.7% formaldehyde, washed with PBS for 5 mins, and mounted on glass slides with DPX. The slides were let dry at room temperature and sealed with nail polish. Cells were visualized at 63X oil objective in a Leica TCS SP8 confocal microscope.

### Degradation assay

Yeast cells were transformed with p315GAL-80QT ataxin-3 and p426CUP-PJA1 plasmids and the tranformants were selected with synthetic drop out media without leucine and uracil (SD-Leu-Ura). The transformed cells were grown overnight in synthetic dropout media with 2% raffinose. PJA1 expression was promoted by addition of 300μM of copper. After 2 hours of PJA1 expression, 2% galactose was added for induction of ataxin-3 protein. Ataxin-3 induction was blocked after 2 hours by addition of 2% glucose, and then chased for further 8 hours. Cells were harvested at regular time intervals.

For cycloheximide chase analysis of 80QT ataxin-3 in PJA1 silenced condition, Neuro2A (N2A) cells were transfected with scrambled and smartpool siRNA specific for PJA1. 6 hours later, the cells were seeded freshly. 18 hours after siRNA transfection, cells were transfected with pEGFP-N1-80QT ataxin-3. Cycloheximide (10μg/ml) was added to the cells to impede translation and chased further for 10 hours with harvesting the cells at regular time intervals.

### Spot assay

Yeast cells transformed with p315GAL-80QT-ataxin-3 and p426CUP-PJA1 plasmids were grown overnight in synthetic dropout media without leucine and uracil containing 2% raffinose. 300μM copper was added to induce PJA1 for 2 hours followed by addition of 2% galactose for induction of ataxin-3. Cells were grown until OD_600_ at 0.5. The cultures were diluted serially and spotted onto YP plate in presence of either 2% galactose with 100μM copper or 2% glucose media. The plates were kept at 30°C for 2 days.

### Ubiquitination assay

Neuro2A cells were transfected with PJA1 specific siRNA and shRNA. 6 hours after transfection, the cells were seeded freshly for the next set of transfection. 18 hours after siRNA transfection, cells were transfected with pCMV-HA-Ubiquitin and pEGFP-N1-80QT. Autophagy inhibitor 3-MA or proteasome inhibitor lactacystin was added to the cells 6 hours before harvesting the cells. The cells were then lysed with cell lysis buffer (10 mM Tris-HCl, pH 8.0, 150 mM NaCl, 2% SDS, 1mM Phenylmethylsulfonyl fluoride with protease inhibitor cocktail). Protein was estimated by BCA method. The lysates were diluted 10 times with IP Dilution Buffer (10 mM Tris-HCl, pH 8.0, 150 mM NaCl, 2 mM EDTA, 1% Triton); anti-GFP antibody was added to the lysate and the immunocomplex was incubated at 4°C overnight with gentle shaking. The next day, protein A-sepharose (GE Healthcare) beads were added to the cell lysate-antibody mixture and incubated at 4°C for 2 hours. The resin was then washed three times with wash buffer (10 mM Tris-HCl, pH 8.0, 1 M NaCl, 1mM EDTA, 1% NP-40) and boiled with 2X SDS loading buffer for immunoblot analysis.

### Reverse transcriptase PCR analysis and semi-quantitative real time PCR analysis

Neuro2A cells were transfected with normal and expanded polyglutamine protein. After 72 hours of transfection cells were harvested and total RNA was isolated with TRIzol reagent (Qiagen). cDNA was synthesized by using RT-PCR kit (Life Technologies). The real time PCR was performed by using SYBR green super mix (Applied Biosystems) using an iCycler iQ real-time Thermocycler Detection System (Applied Biosystems). The RT-PCR analysis conditions were the same for both PJA1 and GAPDH. The following primers were used for detection of PJA1; Forward 5′-GCAATGGCAGTGGTTATCCT-3′ and Reverse 5′- ACATCCCAGGCTGTATGAGC-3′. GAPDH was used as control; Forward 5′- ACCCAGAAGACTGTGGATGG- 3′; and Reverse, 5′-CACATTGGGGGTAGGAACAC-3′. All reactions were carried out in triplicate with negative controls lacking the template DNA.

### RNA interference

Smartpool small interfering RNA (siRNA) of mouse PJA1 were purchased from Dharmacon. Short hairpin RNA (shRNA) targeted to mouse PJA1 was generated using the pLKO.1 shRNA protocol. The primers used for generating shRNA of PJA1 are: Forward 5′- CCGGCACCGACGATTACTACCGATACTCTCGAGAGTATCGGTAGTAATCGTCGGTGTTTTTG and Reverse 5′- AATTCAAAAACACCGACGATTACTACCGATACTCTCGAGAGTATCGGTAGTAATCGTCGGTG. pEGFP-N1-20QT/80QT/130QF ATXN3 and pDsRed-C1-83Q HTT expressing cells were transiently transfected with siRNA, shRNA of PJA1 and control siRNA according to the manufacturer’s protocol.

### Drosophila stocks and husbandry

Fly stocks and crosses were maintained at standard condition (25°C). Rh1-GAL4 was employed for overexpressing various transgenes (Xiong *et al.*, 2012). An UAST-PJA1 transgenic fly was generated. PJA1 was digested from pCMV-HA-PJA1 construct using Not1 and Kpn1 specific forward and reverse primers respectively and cloned into pUAST plasmid vector (Brand and Perrimon, 1993). The plasmid clones were sent to CCAMP, Bangalore, sub-cloned into pUAST-attB, then used for microinjection and targeted integration into the third chromosome. Transgenic flies were scored based on eye colour marker, and the stocks were subsequently balanced. Rh1-GAL4 was used to drive the transgene and the expression of PJA1 was confirmed by Western blot analysis. UAS-Hsap\MJD.tr-Q78 flies harbouring the previously reported truncated ataxin-3 construct was obtained from Bloomington Stock Centre (#8150). Genetic crosses were carried out to make stable stocks by combining this construct with UAST-PJA1 and UAS dicer2 (Bloomington #24644) respectively. Experiments were carried out by crossing Rh1 GAL4 flies to the respective genotypes. The progeny flies were sorted for the appropriate genotypes and were maintained at 25^°^C by regularly transferring to a new vial of fly food. Retinal histology was carried out on 8-10 and 35-40 day old flies.

### Retinal histology

Adult fly heads were fixed in 3.7 % formaldehyde for 60-90 mins in ice. They were next treated with a gradient of sucrose solutions, and subsequently placed in 30% sucrose overnight. The next day 5μm frontal retinal sections were obtained using microtome (Leica). The sections were post fixed with 3.7% formaldehyde and stained with hematoxylin and eosin and observed under microscope at 20X (Leica). At least 10 animals were examined for each genotype.

### Immunofluorescence of retinal sections

The retinal cryosections were fixed in 0.5% formaldehyde in 1X PBS for 30 mins, followed by thorough rinses in 1X PBS. The slides were washed in 0.05% Triton X-100, and then blocked in 3% BSA for an hour. The slides were next incubated in the corresponding primary antibodies overnight. The next day, the slides were washed, and again incubated in Goat-anti-rabbit and Goat-anti-mouse antibodies conjugated to TRITC and Alexa-Fluor 488 respectively for one hour. This is followed by PBS washes, air-drying the slides and mounting with DPX mountant. The slides were viewed at 20X under confocal microscope.

## ACKNOWLEDGEMENTS

We thank Dr. Nitai P. Bhattacharya for his generous amount of input for this study. We are thankful to Dr. Oliver Stork, Dr. Debashis Mukhopadhyay and Dr. Nihar R. Jana for gifting the PJA1, huntingtin and ataxin-3 constructs respectively. We thank the Bloomington Drosophila Stock Center for various *Drosophila* stocks and CCAMP Bangalore, India for generating the UAST-PJA1 transgenic flies. We also thank Dr. Rakesh Mishra, Dr. V. Bharathi, Vijayishwer Jamwal and Jayashish Ghosh for their help with cryosectioning; Dr. Sahana Mitra, Dr. Joydeep Roy, Nilanjan Gayen, Dr. Soumita Mukherjee, Somesh Roy, Madhuparna Chakraborty and Upama Choudhury for their helpful discussions about the work. This work was supported by Bose Institute Intramural fund.

## CONFLICT OF INTEREST

The authors declare no conflict of interest.

## ABBREVIATIONS

PQC: Protein quality control
SCA3: Spinocerebellar ataxia Type 3
HD: Huntington’s disease
PD: Parkinson’s disease
AD: Alzheimer’s disease
PolyQ: Polyglutamine
CHIP: Carboxy-terminus of Hsp70-interacting protein
PJA1: Praja1

## Notes

### Competing Interest Statement

The authors have declared no competing interest.

