## Supplemental File for "Praja1 ubiquitin ligase facilitates degradation of polyglutamine proteins and suppresses polyglutamine-mediated toxicity"

Baijayanti Ghosh[a], Susnata Karmakar[b], Mohit Prasad[b] and Atin K. Mandal[a#]
[a]Division of Molecular Medicine, Bose Institute, Kolkata, India;
[b]Department of Biological Sciences, Indian Institute of Science Education and Research Kolkata, Mohanpur Campus, Mohanpur, Nadia, West Bengal, India

**Supplemental Material**

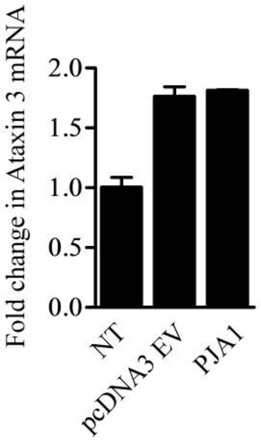


FIGURE S1. ATXN3 transcript levels were unaltered upon over-expression of PJA1. PJA1 along with its empty vector pCDNA3 were transfected in HEK293T cells. 24 hours after transfection, total RNA was isolated from the cells and subjected to quantitative real-time PCR. Data was collected from two separate experiments were normalised to the levels of GAPDH.

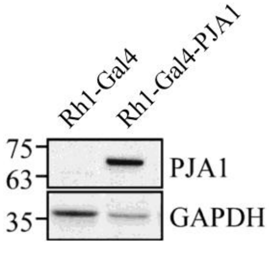


FIGURE S2. Expression of PJA1 in transgenic Rh1-GAL4-PJA1 *Drosophila*. Rh1-GAL4-PJA1 flies were processed for immunoblotting with anti-PJA1 and anti-GAPDH antibodies.

TABLE 1. Plasmids used in this study

| Plasmid | Vector | Description | Source |
| --- | --- | --- | --- |
| 1. Atxin-3 80QT | pEGFP-N1 | GFP-tagged expression plasmid, CMV promoter | Dr. Nihar R. Jana |
| 2. Ataxin-3 20QT | pEGFP-N1 | GFP-tagged expression plasmid, CMV promoter | Dr. Nihar R. Jana |
| 3. Ataxin-3 130QF | pEGFP-N1 | GFP-tagged expression plasmid, CMV promoter | Dr. Nihar R. Jana |
| 4. Ataxin-3 20QF | pEGFP-N1 | GFP-tagged expression plasmid, CMV promoter | Dr. Nihar R. Jana |
| 5. Ataxin-3 20QT | p315GAL | GFP-tagged expression plasmid, Gal promoter, (BamH1+ Not1) | This study |
| 6. Ataxin-3 80QT | P315GAL | GFP-tagged expression plasmid, Gal promoter, (BamH1+Not1) | This study |
| 7. Htt 16Q | pEGFP-C1 | GFP-tagged expression plasmid, CMV promoter | Dr. Debashis Mukhopadhyay |
| 8. Htt 83Q | pDsRed | DsRed-tagged expression plasmid, CMV promoter | Dr. Debashis Mukhopadhyay |
| 9. PJA1 | pCMV- HA | HA-tagged expression plasmid, CMV promoter | Dr. Oliver Stork |
| 10. PJA1 | pDsRed | DsRed-tagged expression plasmid, CMV promoter, (Xho1+EcoR1) | This study |
| 11. PJA1 | pEGFP-C1 | GFP-tagged expression plasmid, CMV promoter | Dr. Oliver Stork |
| 12. PJA1 | pUAST-attB | (Not1+Kpn1) | CCAMP |
| 13. PJA1 | p426CUP | HA-tagged expression plasmid, CUP promoter, (HindIII+Xho1) | This study |
| 14. H553S PJA1 | pCMV-HA | HA-tagged expression plasmid, CMV promoter | This study |
| 15. ΔRING PJA1 | pCDNA3.1 | HA-tagged expression plasmid, CMV promoter, (HindIII+Not1) | This study |
| 16. shPJA1 | PLKO | (Age1+EcoR1) | This study |
| 17. EZH2 | pCMV-HA | HA-tagged expression plasmid, CMV promoter | Addgene |
